# Advantages of using Empirical Mode Decomposition and Hilbert Transformation for Delineating Resting State Functional Brain Networks

**DOI:** 10.64898/2026.04.23.719165

**Authors:** Taranjit Kaur, Sandeep Yadav, Neeraj Jain

## Abstract

The goal of the resting-state functional connectivity studies is to determine the inherent dynamics of the brain networks while the body is at rest. These networks get differentially activated when the brain is involved in various tasks such as processing of sensory inputs, initiating motor activities, or various cognitive tasks. Resting state functional connectivity networks are commonly revealed by determining Pearson Correlation Coefficients of the Blood Oxygenation Level Dependent (BOLD) signals collected from different brain regions using functional Magnetic Resonance Imaging (fMRI) while the subject is not actively performing any task. However, the functional connectivity thus determined does not correlate well with the known structural connectivity between different brain regions. Here, we used Empirical Mode decomposition (EMD), followed by Hilbert Transformation (HT), to determine the resting state functional connectivity in the human brains. We show the advantage of using this EMD-HT method using somatomotor network as an example. We show that the time series data decomposed by this method improves correlation of the derived functional connectivity with the known structural connectivity (especially for low -TR fMRI data) as compared to the methods commonly used.

## 1. INTRODUCTION

The functional magnetic resonance imaging (fMRI) of the brain measures Blood Oxygenation Level Dependent (BOLD) signals, which correlate with the level of activities in different areas of the brain. BOLD are pre-processed and used to determine temporal coherence in different brain regions to reveal the functional connectivity (FC) networks composed of coactive areas of the brain. Functional connectivity networks revealed while the brain is not actively involved in any sensory, motor, or cognitive task, reveals resting-state brain networks (RSN)(Biswal et al., 1995)(Thomas et al., 2021) (Wei et al., 2024). The widely used method to determine FC between different areas of the brain involves applying a band-pass filter (BPF) to the time series data and then computing the Pearson correlation between the Regions of Interest (ROI) pairs. Higher values of the temporal correlations between brain regions suggest stronger FC between those brain regions. This commonly used method makes underlying assumption that the BOLD signals are stationary and exhibit linearity. However, the BOLD signals exhibit complex dynamics that vary over time, having both non-linearity and non-stationarity. Therefore, the BPF method leads to incomplete network connectivity inferences.

Moreover, the resting state-Functional Connectivity (rsFC) thus determined does not correlate well with the known structural connectivity (SC) (Thomas et al., 2021)(Honey et al., 2009).

Empirical Mode Decomposition (EMD) based method is another tool to determine FC (Zhang et al., 2015) (Szakál, 2016) (Yuen et al., 2019)(Das et al., 2020). EMD divides the time series data into a set of components called Intrinsic Mode Functions (IMFs) that represent the oscillatory behaviour at different time scales (Huang et al., 1998). Therefore, EMD-based methods could be better in deciphering the complex dynamics of the BOLD fMRI signal. Further, EMD when combined with Hilbert transformation, gives an estimate of the temporal evolution of the phase and frequency. EMD has been used for rs-fMRI data, including multivariate empirical mode decomposition (MEMD) and Variational Mode Decomposition (VMD) to reveal FC of the visual and motor cortices ((Szakál, 2016), default mode networks (DMN) (Zhang et al., 2015), and topology of resting state brain wide network (Yuen et al., 2019).

We considered somatomotor network, to compare rsFC by two different methods. One, using BPF on time series data from different brain regions, i.e., the BPF method. Two we used the same time series data decomposed by EMD into IMFs and further analyzed by Hilbert Transformation (HT), i.e., the EMD-HT method. We compared the FC thus determined using the two different methods with the known SC. We show that EMD, followed by selection of meaningful components based on mean instantaneous frequency computed via HT, gives better correlation with SC.

## 2. MATERIALS AND METHODS

We used fMRI data from three different sources acquired using different parameters to establish the robustness of our inferences. Dataset 1 was collected by us. Dataset 2 was from an online dataset made available as part of the Human Connectome Project (Fallon et al. (Fallon et al., 2020)), and Dataset 3 was from the Population Imaging of Psychology 2 (PIOP2) dataset available as a part of the Amsterdam open MRI collection(Snoek et al., 2021).

We present the details of the data collection procedure for Dataset 1. For other datasets, the procedures are available as described in relevant references. Here, we will highlight only the differences in the acquisition parameters from Dataset 1.

### 2.1 Acquisition of Dataset 1

#### 2.1.1 Subjects and acquisition parameters

We acquired data from ten right-handed subjects (five males and five females) aged 22-39 years. The study participants did not report any history of psychiatric or neurological illness. Informed consent was obtained from the participants. The protocols were approved by the Human Ethics Committee of the National Brain Research Centre, where the data was acquired (Thomas et al., 2021). Other unrelated results from these data were reported earlier (Thomas et al., 2021). Data was acquired using a 3T MR scanner (Phillips, Achieva, Netherlands). The anatomical T1-weighted data was acquired using Repetition Time (TR)=8.4 ms; Field of View (FOV)=250 mm x 230mm x 150mm; 150 slices. The functional scans in resting state were taken with T2-weighted gradient echo sequence with TR=2000 ms; TE=30ms, FOV = 230 mm x 242 mm x 132 mm; slice thickness 4 mm, 33 slices. A total of 205 functional volumes were acquired for each participant who was lying in a supine position in the bore of the magnet. The subjects were instructed to stay still and not to actively think any particular thoughts as far as possible.

#### 2.1.2 Data pre-processing and extraction of time series data from the Regions of Interest (ROIs)

The fMRI data processing was carried out using statistical parametric mapping (SPM12; http://www.fil.ion.ucl.ac.uk/spm) software operating in Matlab 2022b environment (MathWorks Inc., MA, USA). Data pre-processing steps have been described before (Thomas et al., 2021). First, the functional images were corrected for slice time to ensure that all slices in the volume were acquired simultaneously. Second, the volumes were corrected for head motion using a six-parameter ‘rigid body transformation’ to minimize the difference between the first scan (usually the reference scan) and the successive scans. Third, the functional volumes were co-registered with anatomical images. Finally, normalization was done to align the functional data to a standard template (ICBM 2009a Nonlinear Symmetric template) and resampled to 2-mm cubic voxels.

For extraction of the time series data from specific ROI’s, we used the Glasser brain atlas to demarcate the ROI’s(Glasser et al., 2016). The atlas provides parcellation of the brain into three hundred and sixty regions. For each region, the provided mask image was then applied to the pre-processed fMRI data to extract the time series data from each voxel. Before applying the mask, we ensured that the atlas and the functional data were in the same space using Montreal Neurological Institute (MNI) template (Mazziotta et al., 1995). Thereafter, the time series data from each voxel within an ROI were averaged to generate one signal for each of the brain regions. These averaged signals were used to determine the correlations. Since we were interested in the resting-state somatomotor network, the somatosensory cortical areas 1, 2, 3a, and 3b, second somatosensory area (S2), primary motor area 4, medial area 5 (5m), lateral area 5 (5L), dorsal area 6 (6d), ventral area 6 (6v), supplementary motor area (6mp, 6ma), supplementary and Cingular eye field (SCEF), were used as nodes of the network. Thus, data was considered from 13 cortical areas (nodes) per hemisphere, for a total of 26 different nodes from the two hemispheres. The Glasser parcellation subdivides regions 5, 6, and SMA into multiple sub-regions, resulting in several time series per region. Given the spatial proximity of these sub-regions, we averaged the time series data from these regions ignoring the subdivisions to avoid redundancy. The time series data from ROI’s defined as area 5m and area 5L in the Glasser parcellation (Glasser et al., 2016)) were averaged and designated as the signal from area 5; data from area 6d and area 6v were averaged and designated as signal from area 6; data from areas 6mp, 6ma and SCEF was averaged and designated as signal from supplementary motor area (SMA). This reduced the number of nodes from 26 to 18, i.e., 9 nodes per hemisphere. Here, we determined connectivity of the somatomotor network independently for each hemisphere and averaged FC value between each pair of nodes from the two hemispheres.

### 2.2 Dataset 2

Dataset 2, the parcelled time series data of Fallon et al. (Fallon et al., 2020) were generated from the resting state fMRI scans of 100 subjects from the Human Connectome Project (HCP) (Van Essen et al., 2013). The acquisition parameters were TR=720 ms, TE=33.1 ms, slice thickness =2mm, and the number of time points=1200. This dataset is available in pre-processed and pre-parcellated using the Glasser atlas, the same that we used for Dataset 1. Here, we used data of 10 randomly selected subjects.

### 2.3 Dataset 3

Dataset 3 is publicly available resting state fMRI dataset from the Amsterdam open MRI collection (Snoek et al., 2021). This pre-processed PIOP2 dataset was acquired using TR of 2 seconds, the same as Dataset 1. A total of 10 subjects were randomly taken for the analysis. The acquisition parameters were TE=28 ms, slice thickness =0.3 mm, and the number of time points = 240. We have used the pre-processed version of the PIOP2 dataset, which was subsequently parcelled via the Glasser atlas by us to extract the time series from the regions of interest (Snoek et al., 2021).

For further analyses of all the datasets- Dataset 1, 2 and 3, we followed the same methodology outlined below.

### 2.4 Empirical Mode Decomposition (EMD)

Empirical Mode Decomposition (EMD) is a data-driven approach used for the analysis of non-linear and non-stationary signals (Huang et al., 1998). We performed the EMD decomposition process for a signal *x*(*t*) as follows(Zeiler et al., 2010): (1) computed the local maxima and minima, which were denoted by *P*_i_ and *p*_i_ where *i* =1, 2, ….., in *x*(*t*); (2) using cubic spline interpolation method calculated the upper and lower signal envelope: *P*(*t*)*=f_P_*(*P_i_,t*) and *p*(*t*)*=f_p_*(*p_i_, t*); (3) calculated the envelope average as *e*(*t*)=(*P*(*t*)+*p*(*t*))/2; (4) calculated the details as *s*(*t*)=*x*(*t*)-*e*(*t*). We then examined the following properties of *s*(*t*): (A) if *s*(*t*) satisfied the conditions on the number of extremas & zero crossing and the symmetry about local mean, calculated the *i*^th^ IMF as *s*(*t*); i.e., *IMF*(*t*)=*s*(*t*) and change *x*(*t*) with the residual value *r*(t)=*x*(*t*)-*IMF* (*t*): (B) If *s*(*t*) was not an IMF, then *x*(*t*) was replaced with detail value, i.e., *x*(*t*)=*s*(*t*).

Steps (1) to (5) were iteratively executed until *r*(*t*) satisfied a stopping criteria *D* as:

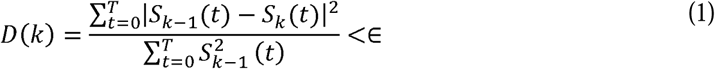

where *k* is the *k*^th^ difference between the signal *x*(*t*) and the envelope *e*(*t*) mean. The parameter ε is a pre-determined constant value governing the stopping of the shifting process. The signal *x*(*t*) was then reconstructed as follows:

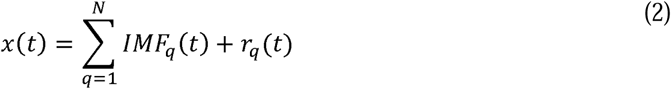

where *N* is the IMF number (satisfying criteria of orthogonality and nearly zero mean) and *r_q_* (*t*) is the ending residue value signifying the low-frequency trend of the signal.

### 2.5 Hilbert Transformation and the Instantaneous Amplitude and Frequency

After extracting the IMF using EMD, we calculated the Hilbert transformation of the extracted IMF as given by the following equation (Huang et al., 1998):

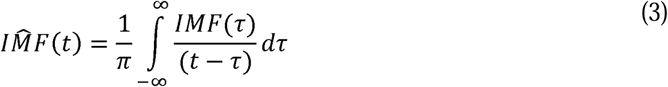

where 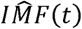 is the Hilbert transformation of the IMF. After Hilbert transformation, we calculated the analytic signal *y*(*t*) as:

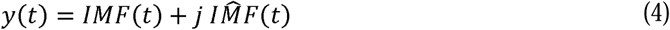

This gave a complex valued signal where the real part denotes the IMF, and the imaginary part signifies the Hilbert transformation. Few other parameters were computed using equations (3) and (4). These were Instantaneous Amplitude, A(t); Instantaneous Phase, *φ*(t); and Instantaneous Frequency, *f*(*t*), mathematically represented using the equations below:

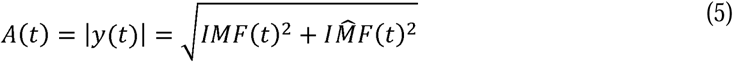

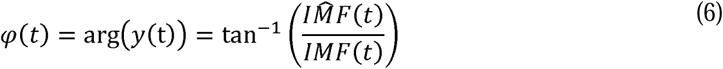

Taking the derivative of the instantaneous phase and dividing it by a constant of 2π, we obtained the instantaneous frequency i.e. the frequency content of the signal at each time instant.

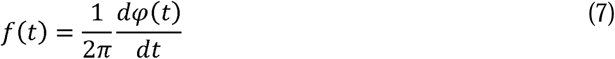

An example of the above framework for Dataset 1 is shown in Fig. 1-4. As described above, the fMRI data was collected from 20 hemispheres of 10 subjects, and time series was extracted from 9 nodes of the somatomotor network in each hemisphere (Table 1); i.e. for a total of 18 signals from both the hemispheres.

**Fig. 1.**
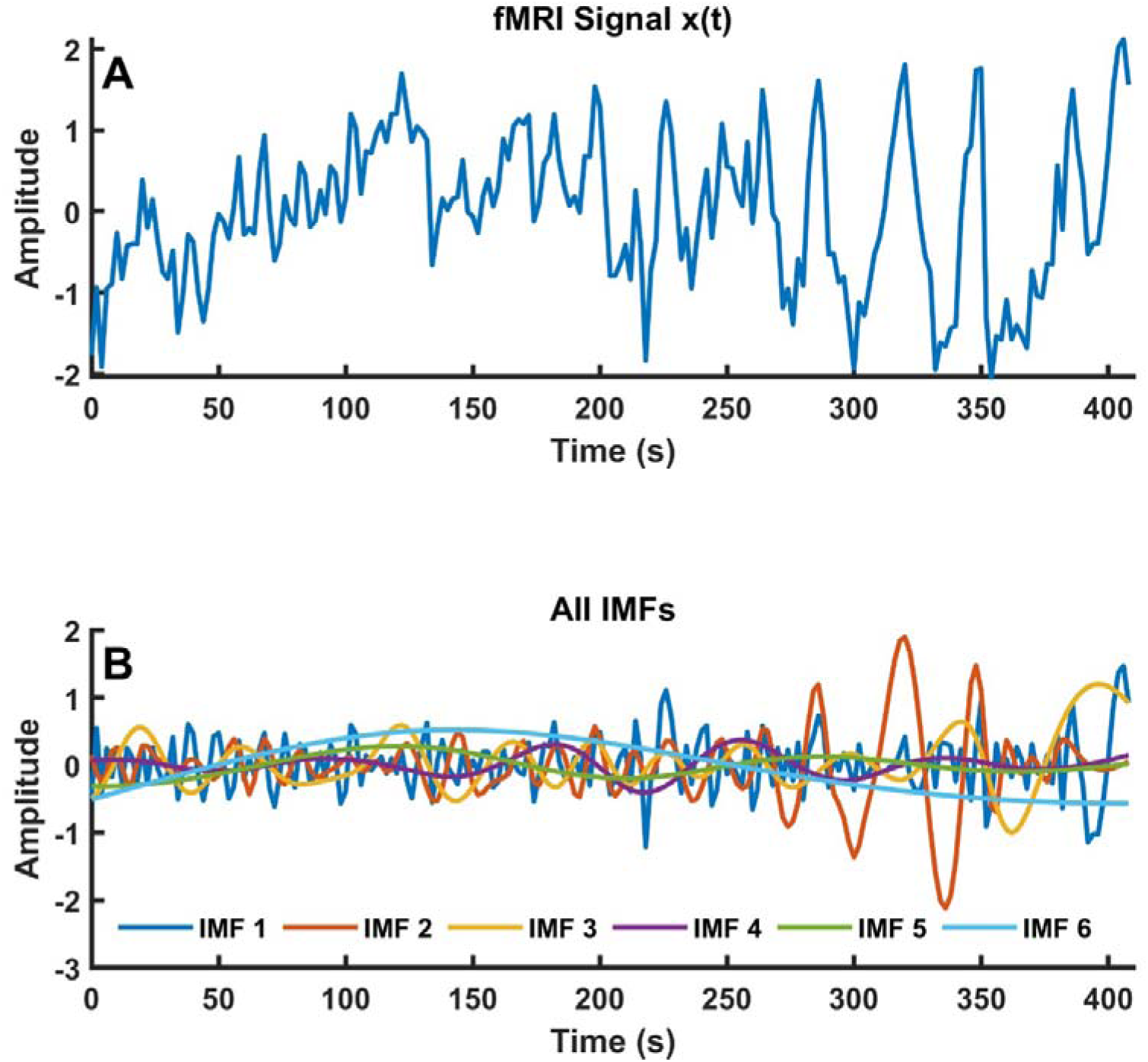
(A) Raw BOLD signal from the right second somatosensory area. (B) Components 1-6 after decomposing by EMD (IMF’s).

**Table 1.** List of the nodes of the somatomotor network.

| Brain area | Node name |
| --- | --- |
| Area 1 | A1 |
| Area 2 | A2 |
| Area 3a | A3a |
| Primary Somatosensory Cortex | A3b |
| Primary Motor Cortex | A4 |
| Area 5 | A5 |
| Supplementary Motor Area | SMA |
| Area 6 | A6 |
| Secondary Somatosensory Area | AS2 |

Fig. 1A shows representative raw time series signal for the secondary somatosensory cortex, S2, from the right hemisphere of one subject. Here, applying EMD to the time series data resulted in six IMF’s (IMF1-6), representing the intrinsic oscillatory components across various frequency scales (Fig. 1B). From different cortical areas (nodes), the signal decomposed in 5-7 components. After applying the Hilbert transformation to the decomposed IMF’s, the instantaneous frequency (IF) for each IMF was obtained (Fig. 2A-F). Using the mean value of the frequency from each IF plot (see the dashed red lines), the IMFs lying in the frequency range of 0.01 to 0.12 Hz (Fig. 3) were used to generate a combined signal merging those IMFs (Fig. 4). The Pearson’s correlation was computed using this combined signal for all the node pairs and the connectivity thus evolved was referred as rsFC_EMD-HT_.

**Fig. 2.**
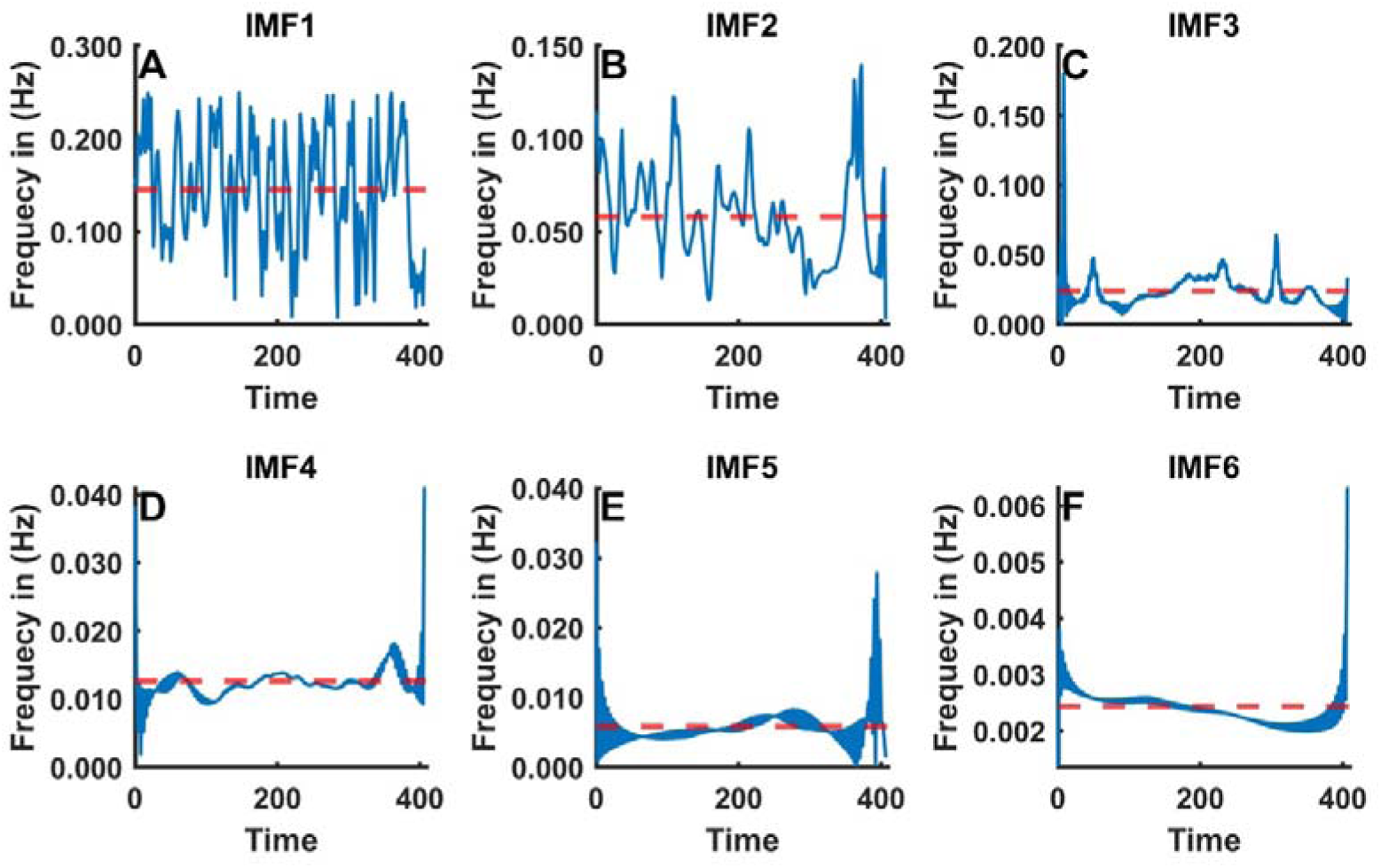
(A-F) Instantaneous frequency plots (blue curves) for all IMFs. The mean value for each IMF is shown by the horizontal dashed red line.

**Fig. 3.**
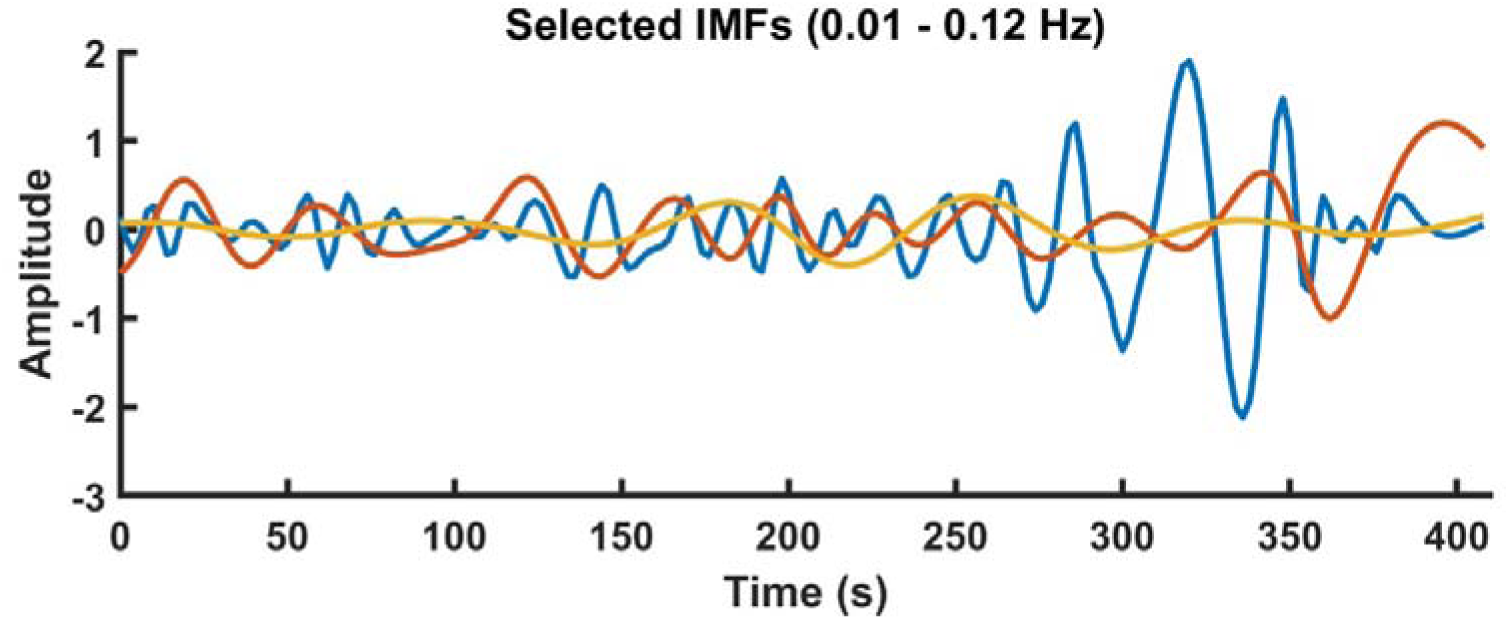
The selected IMFs between 0.01 Hz and 0.12 Hz.

**Fig. 4.**
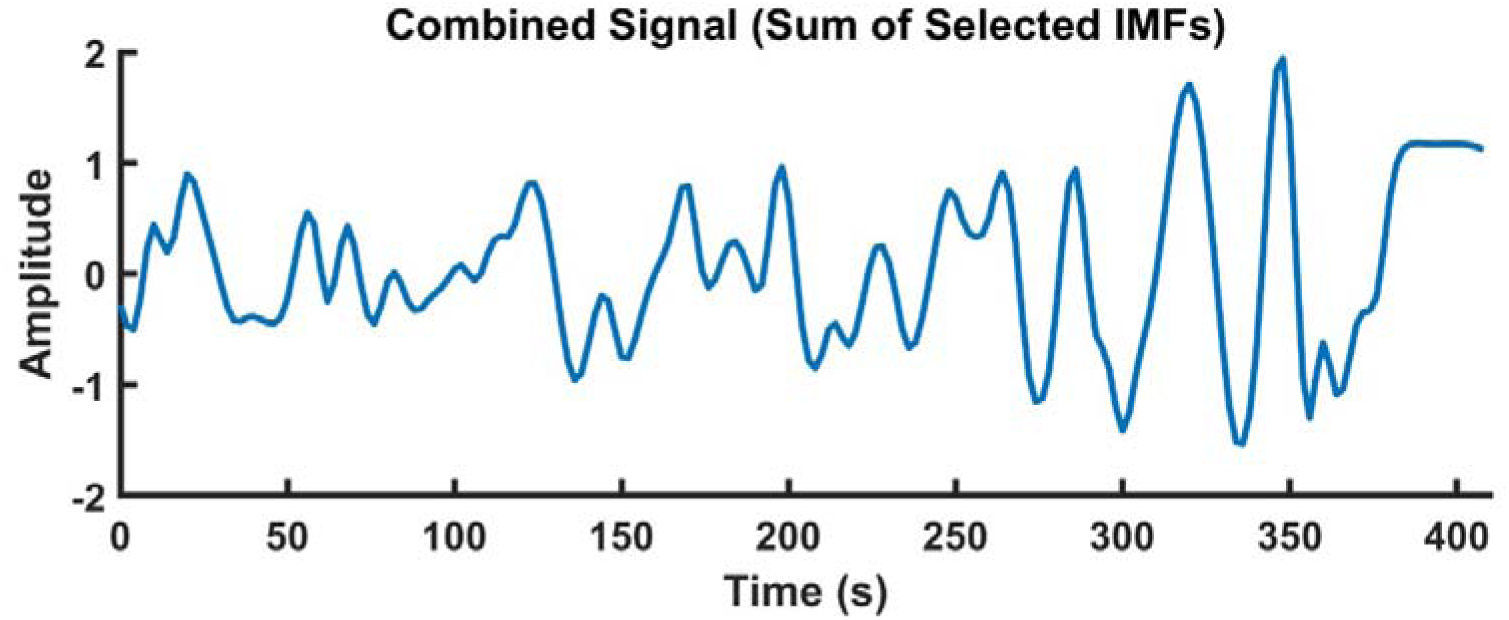
Combined Signal after merging the selected IMFs shown in Fig. 3.

### 2.6 rsFC Computation and comparison with SC

The raw signal was band pass filtered in the range from 0.01 to 0.12 Hz and thereafter Pearson’s correlation coefficients between the ROI pairs was determined. The rsFC in this case is referred to as rsFC_BPF_. The frequency ranges for BPF filtering were chosen to capture the low-frequency fluctuations present in the BOLD data, and at the same time avoiding physiological artifacts from respiratory and cardiac signals(Wang et al., 2022; Wu et al., 2008).

Both rsFC_BPF_ and rsFC_EMD-HT_ were thresholded and compared with the SC proposed by Felleman and Van Essen (Felleman and Van Essen, 1991) (structural connectogram is shown in Fig. 5). Although the SC originates from the tracer studies in macaques, it has been widely used as an anatomical reference model in human neuroimaging studies(Van Essen et al., 2016)(Van Essen et al., 1998). Comparative studies using diffusion MRI based tractography have shown that brain-wide wiring in macaques and human brain is largely similar(Goulas et al., 2014).

**Fig. 5.**
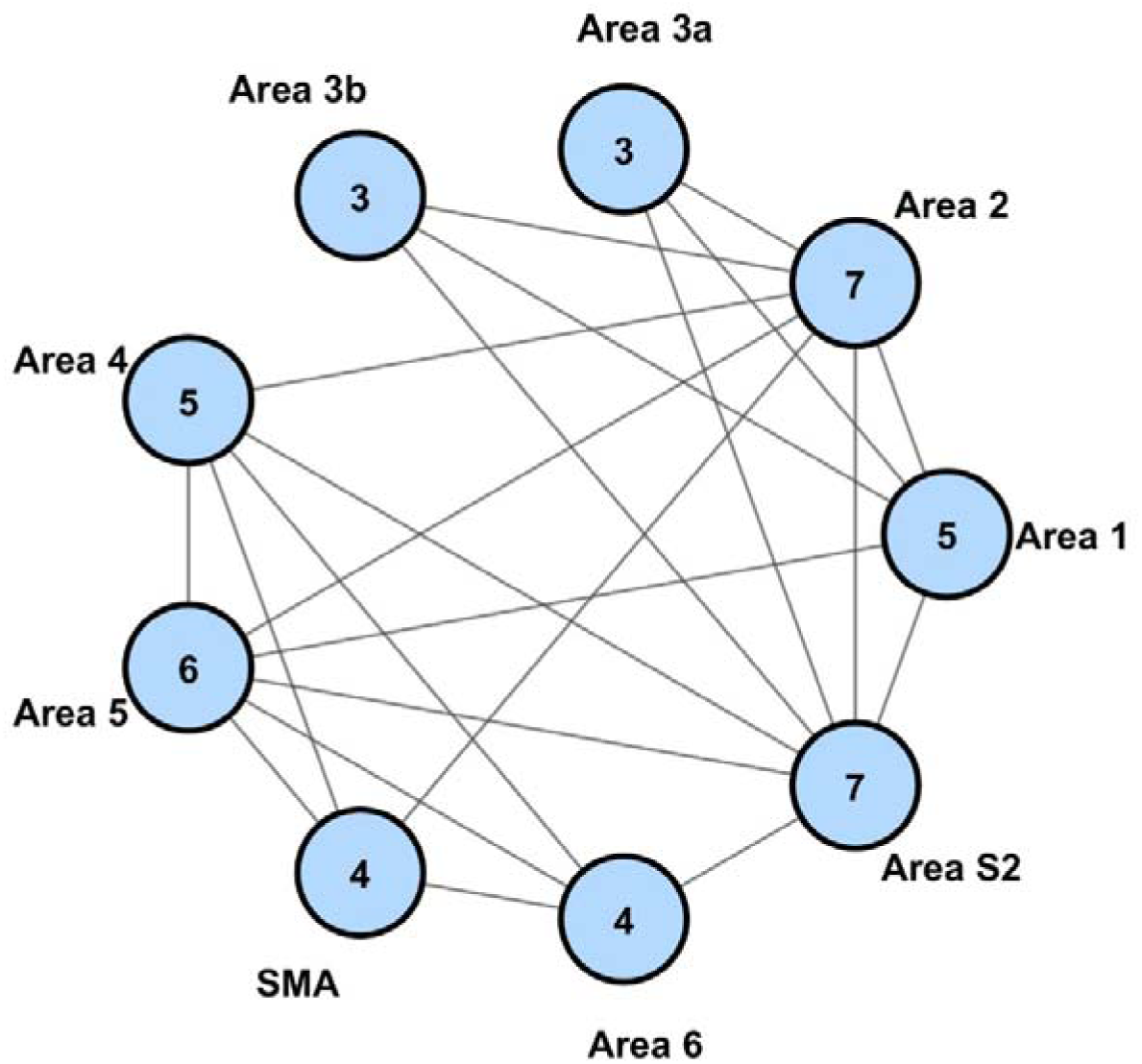
Structural connectome based on Fellman and Van Essen (Felleman and Van Essen, 1991).

### 2.7 Identifying the optimal threshold

For identifying the optimal threshold, three FC measures, i.e., global efficiency (Latora and Marchiori, 2001), modularity(Newman, 2004), and clustering coefficient (Watts and Strogatz, 1998) were computed for a range of the correlation thresholds. As these measures have different numerical ranges, each FC measure was independently normalized in the range from [0,1]. These normalized values were then summed up to form a composite metric (Fig. 6 (D-E), (I-J), and (N-O)). Thereafter we used a triangle method to identify the elbow point (Satopää et al., 2011).

**Fig. 6.**
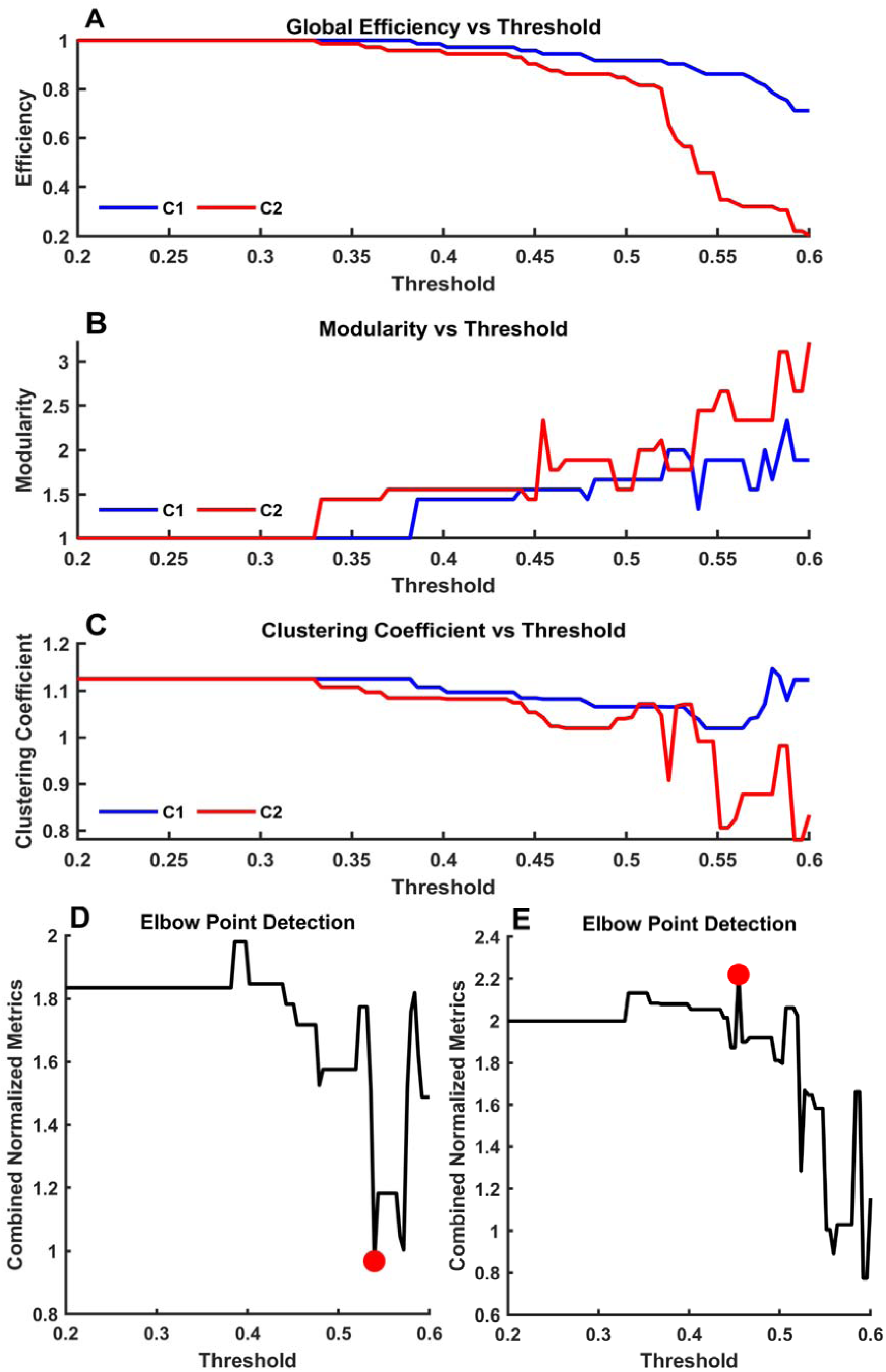
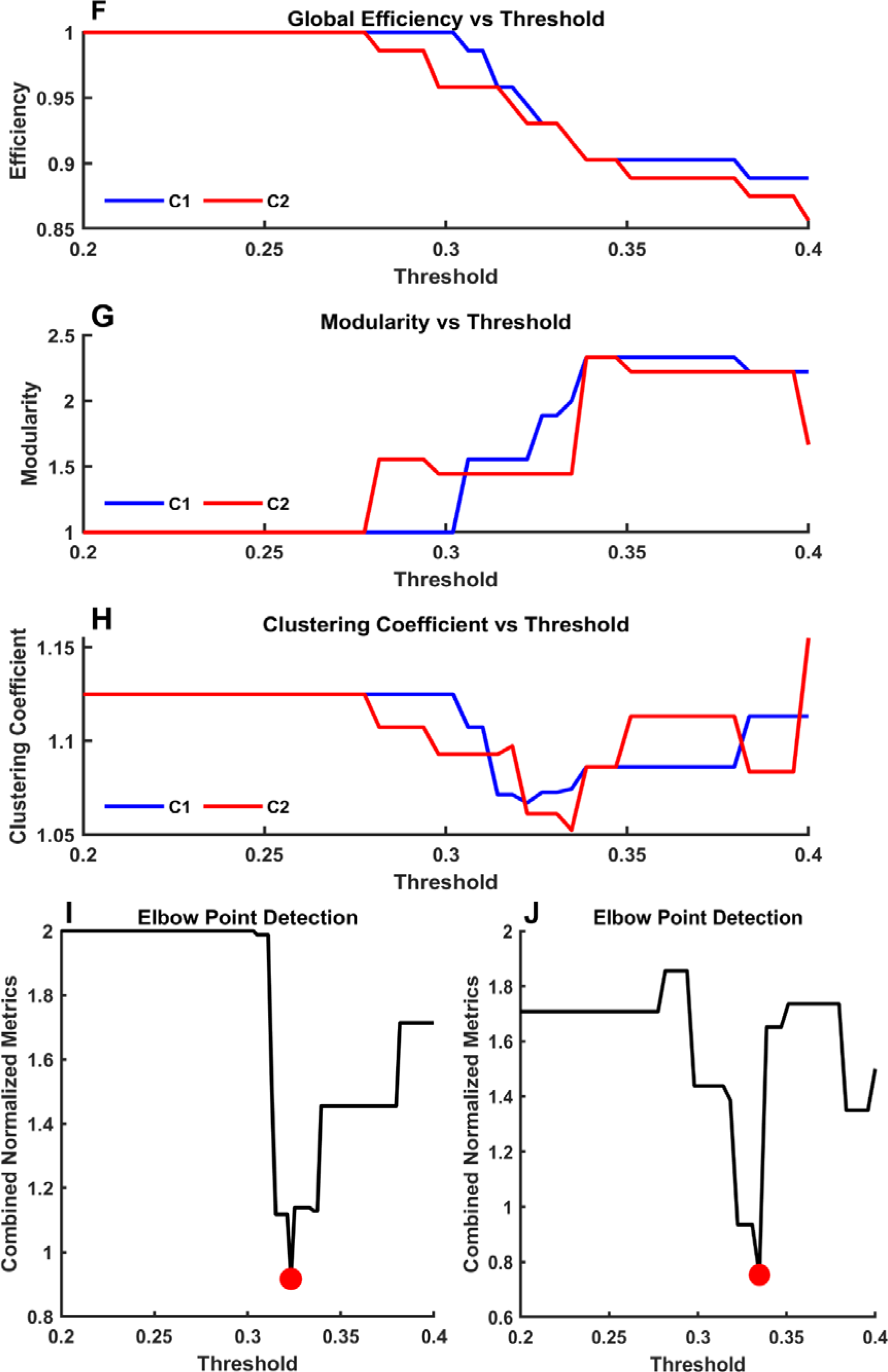
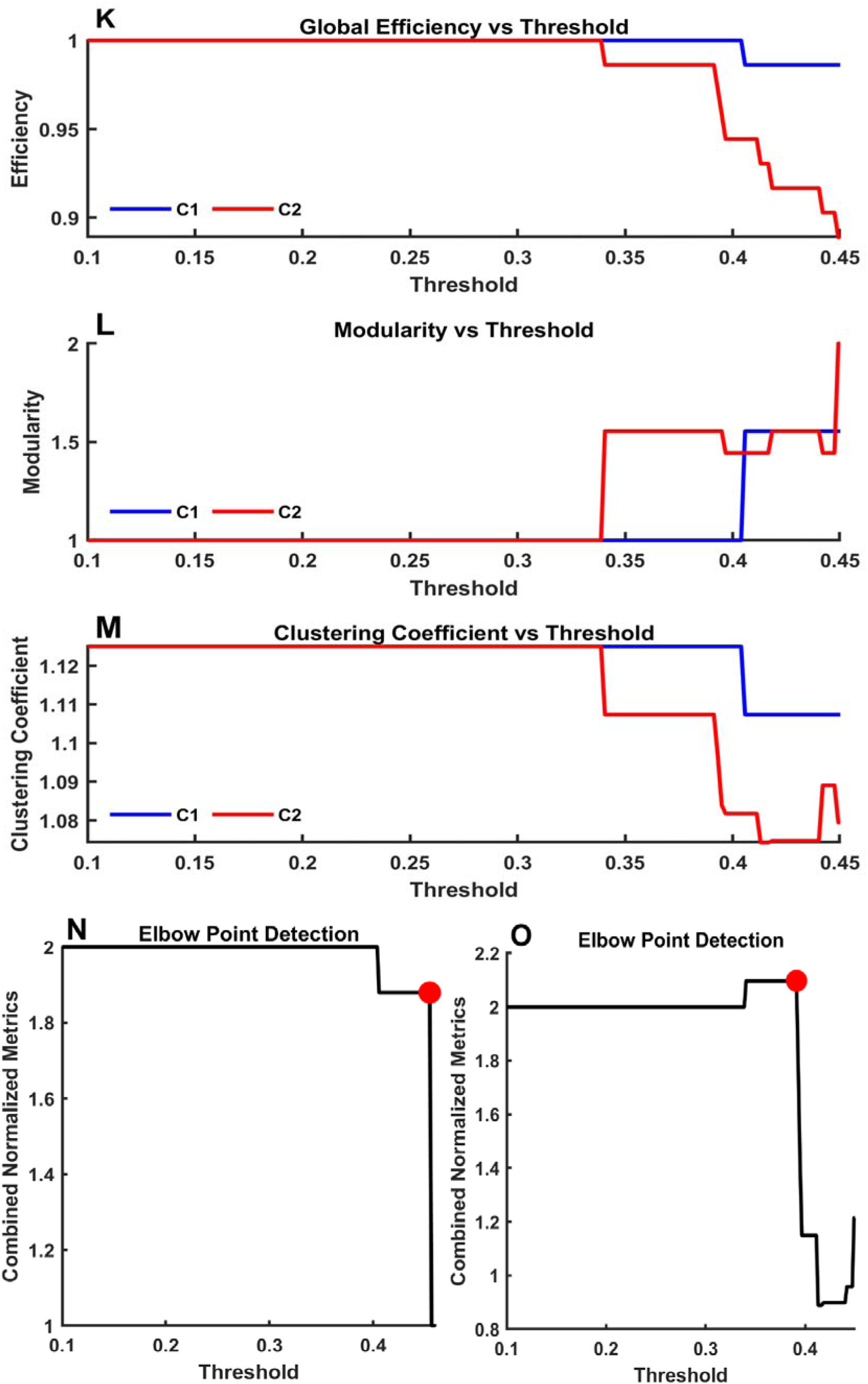
Optimal value of the threshold for the three datasets: (A-C) Variation in the network measures for Dataset 1, (D-E) Elbow point for Dataset 1, (F-H) Variation in the network measures for Dataset 2, (I-J) Elbow point for Dataset 2, (K-M) Variation in the network measures for Dataset 3, (N-O) Elbow point for Dataset 3.

This elbow point (corresponding to the optimal threshold) is a location on the curve that has the maximum perpendicular distance from the line connecting extreme points of the composite metric curve. As the metric curve and elbow point calculations make use of the correlation values having different statistical distributions, the optimal threshold was separately calculated for rsFC_BPF_ and rsFC_EMD-HT_. The obtained threshold values were 0.54, 0.45; 0.32, 0.33, and 0.44, 0.39 for three datasets using the rsFC_BPF_ and rsFC_EMD-HT_.

The BPF-derived network exhibits higher global efficiency, a higher clustering coefficient, but a lower modularity (shown as C1 in blue colour – Fig. 6 (A-C), (F-H), and (K-M)). This implies that in the BPF method, the brain regions are strongly connected both locally and across the entire network, indicating an information flow between distant brain areas. In contrast, the EMD-HT method showed higher modularity (shown as C2 in red- Fig. 6 (A-C), (F-H), and (K-M), i.e., an organization into more distinct communities/sub-modules with stronger functional connections within submodules, For example, primary somatosensory areas (Area 1, Area 2, Area 3a, Area 3b), motor planning/execution areas (Area 4, Area 6, SMA) and higher order somatosensory and associative regions (Area S2, Area 5) and weaker functional associations between sub-modules. Additionally, the identified elbow points for C2, i.e., 0.45, 0.33, and 0.39, encompass both stronger and weaker connections, emphasising a modular dynamic organisation of brain activity closer to the brain’s naturally functioning networks.

We binarized the rsFC _BPF_ and rsFC_EMD-HT_ using the obtained optimal threshold point and assumed that rsFC greater than that point indicate a stronger functional connection to the node.

It has been shown that structurally connected regions show stronger functional connectivity during rest(Hagmann et al., 2008). We defined the terms True Positive (TP), True Negative (TN), False Negative (FN) and False Positive (FP) as follows: TP- both structural as well as functional connectivity present, TN- no structural and functional connectivity present, FP- functional connectivity present but no known structural connectivity, and FN- structural connectivity present without any observed functional connectivity.

We computed the values of TP, TN, FP, and FN and report the performance comparison of the BPF method and the EMD method in the results section.

## 3. RESULTS

We first present results from our own dataset (Dataset 1) with a detailed description of the results. Subsequently, results from Dataset 2 and Dataset 3 are described in brief.

### 3.1 Dataset 1

For Dataset 1, Fig. 7 shows FC matrices with temporal correlations for the BPF method (Fig. 7A, rsFC_BPF_) and the EMD method (Fig. 7B, rsFC_EMD-HT_).

**Fig. 7.**
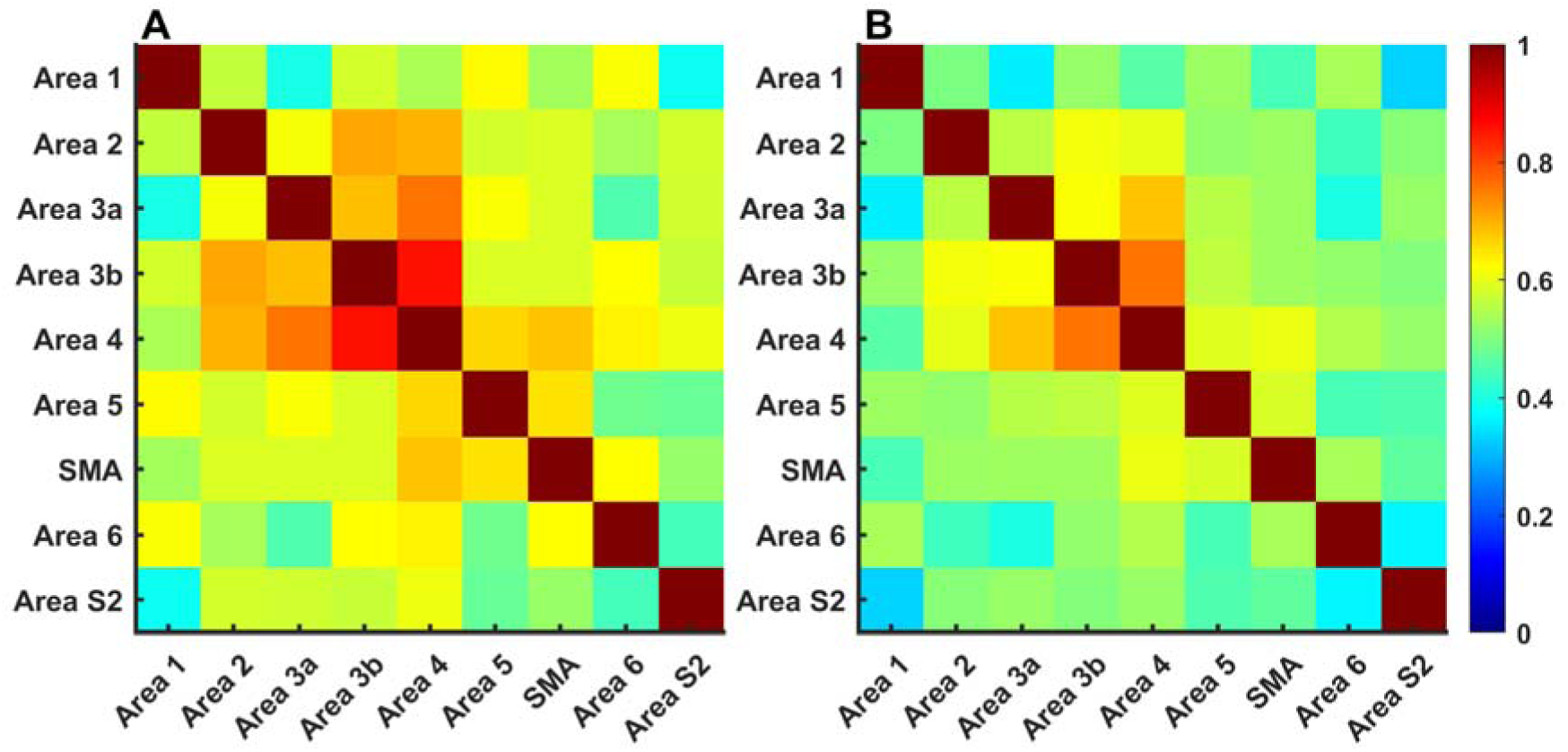
FC matrix for the somatomotor network for Dataset 1 using (A) the BPF method, (B) the EMD method. ROIs are marked along the x-axis and y-axis; the colours show the strength of the FC (Pearson correlation coefficient values) as per the colour scale on the right, which is the same for both ‘A’ and ‘B’. The correlation coefficient was normalised in the range of 0 to 1.

Structure-function correspondence analysis reveals that the functional connections obtained using the EMD method exhibit a higher correspondence with SC. For example, area 5 is structurally connected to several regions, including area 1, area 2, area 4, SMA, area 6, and area S2, but not with area 3a and area 3b (Fellman and Van Hessen (Felleman and Van Essen, 1991)). The connectivity strengths between area 5 and the structurally linked regions were consistently in the moderate to high range, i.e., 0.48-0.67 for the BPF method, and 0.44-059 for the EMD method (Fig. 7).

In contrast, the FC between area 5 and areas 3a and 3b, which lack SC, was high by both methods (0.62 and 0.59 with the BPF, and 0.55 and 0.56 with the EMD).

Upon thresholding the rsFC_BPF_ and rsFC_EMD-HT_ (at the threshold values shown in Fig.6 (D-E), some structurally valid connections are not preserved in the BPF method. In particular, the connection between area 5 and area S2 is retained using the EMD method, whereas it is discarded in the BPF method.

A quantitative comparison of the BPF and the EMD methods using the confusion matrix-derived performance measures is shown in Table 2.

**Table 2.**
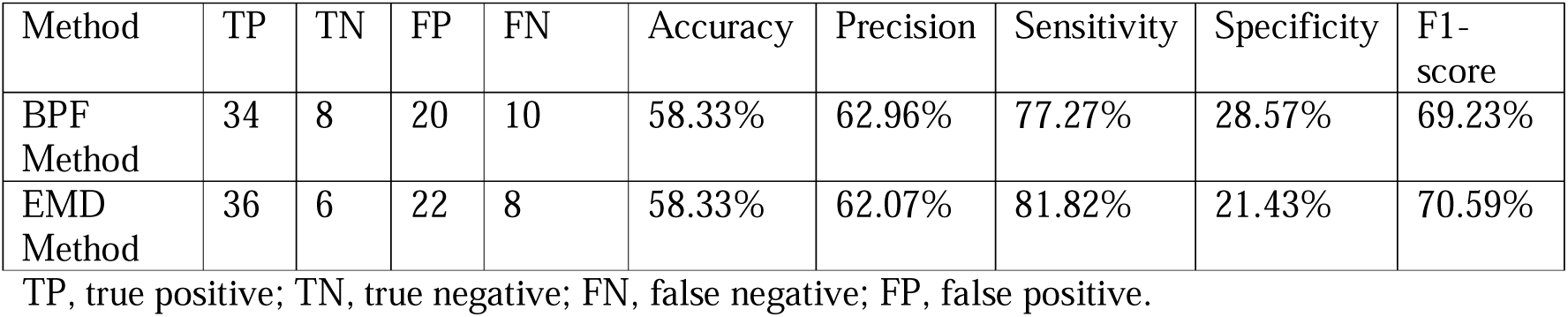
Confusion matrix-derived performance metrics for Dataset 1.

The data show that the sensitivity and F1-score are better with the EMD method. The EMD method yielded accuracy, precision, sensitivity, specificity, and F1-score values of 58.33%, 62.07%, 81.82%, 21.43%, and 70.59%, respectively, whereas the BPF method provided values of 58.33%, 62.96%, 77.27%, 28.57%, and 69.23%, respectively.

To further evaluate the performance of the BPF and EMD methods in extracting SC patterns, we conducted a structural-functional correspondence analysis (Fig. 8) to identify matches and mismatches between the FC obtained using the two methods, i.e., rsFC_BPF_ and rsFC_EMD-HT_, and the direct SC. The connectograms illustrate the two mismatch situations between the functional and the structural networks. One, functional connections not revealed in SC (shown in blue colour, Fig. 8A and 8C), i.e. the functional connections identified from the resting state data (rsFC_BPF_=1 and rsFC_EMD-HT_=1) that lack direct structural evidence (A=0). Two, structural connections not revealed in FC (shown in red colour, Fig. 8B and 8D), i.e. the direct structurally defined pathways (A=1) that do not show corresponding functional connectivity (rsFC_BPF_=0 and rsFC_EMD-HT_=0).

**Fig. 8.**
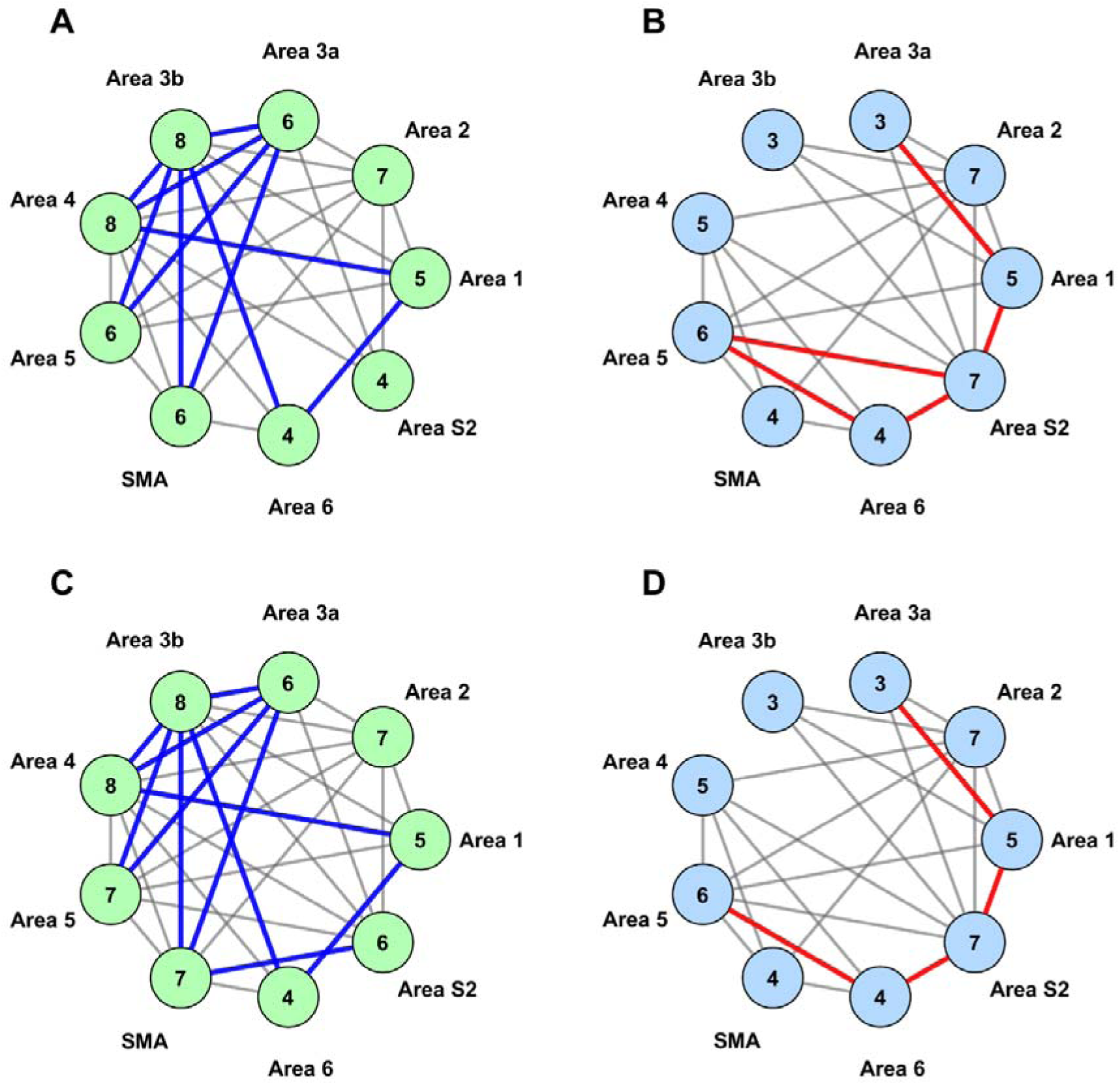
Mismatch between the SC and FC between different nodes of the somatomotor network. Panels (A) and (C) show the group-average and thresholded functional connectograms computed via the BPF and EMD methods. Grey lines show functionally positive pairs survived after thresholding, and the thick blue lines show functional only connections not revealed in SC. Panels (B) and (D) show the corresponding structural connectogram, which is based on the macaque cortical connectivity proposed by Fellman and VanEssen (Felleman and Van Essen, 1991). Grey lines show that structurally positive pairs as per (Felleman and Van Essen, 1991), and the thick red lines show structural connections not revealed in FC. These structural maps serve as a structural reference for the biological basis of the FC.

There were 10 functional connections between areas which have no known SC in BPF and 11 in EMD. The structurally connected areas that did not show any FC were 5 in BPF and 4 in EMD. Thus, EMD showed one extra functional connection, while BPF missed one structural connection, suggesting that rsFC_EMD-HT_ correlated better with the structural connectivity in terms of preserving FC of structurally positive connections.

While analysing mismatches between SC and FC, we also considered the importance of a particular node in the network by calculating the degree centrality (DC) measure, a commonly used measure in the brain network analysis to determine the importance of a particular node in the network (Wang et al., 2010). Any effective approach should reliably identify the key regions with a high DC. In the somatomotor network, regions typically characterised by high DC include the primary somatosensory cortex, secondary somatosensory cortex, primary motor cortex, and supplementary motor areas (SMA). Both the BPF and EMD methods successfully highlighted these areas as highly central within the network, as indicated by the DC values in the connectivity spheres of Fig. 8. The comparable DC values for both the BPF and the IMF method shows robustness of both the techniques in capturing functionally important brain regions in the network.

It is important to note that while the high DC value suggests a region’s functional importance, they do not necessarily indicate a strong structural correspondence. FC studies tend to overestimate the connectivity because they also account for indirect pathways (Honey et al., 2009)

### 3.2 Dataset 2

Dataset 2 was from ten of the subjects from the work by Fallon et al. (Fallon et al., 2020). Fig. 9 shows the FC matrices obtained using the BPF and the EMD methods. Here, as for Dataset 1, the temporal correlations obtained using EMD exhibited better structural correspondence than those obtained using the BPF method for most of the ROI pairs. For example, area 1 has a structural connection to area 2, area 3a, area 3b, area 5, and area S2 and no connection to area 4, area 6, and SMA (Felleman and Van Essen, 1991).

**Fig. 9.**
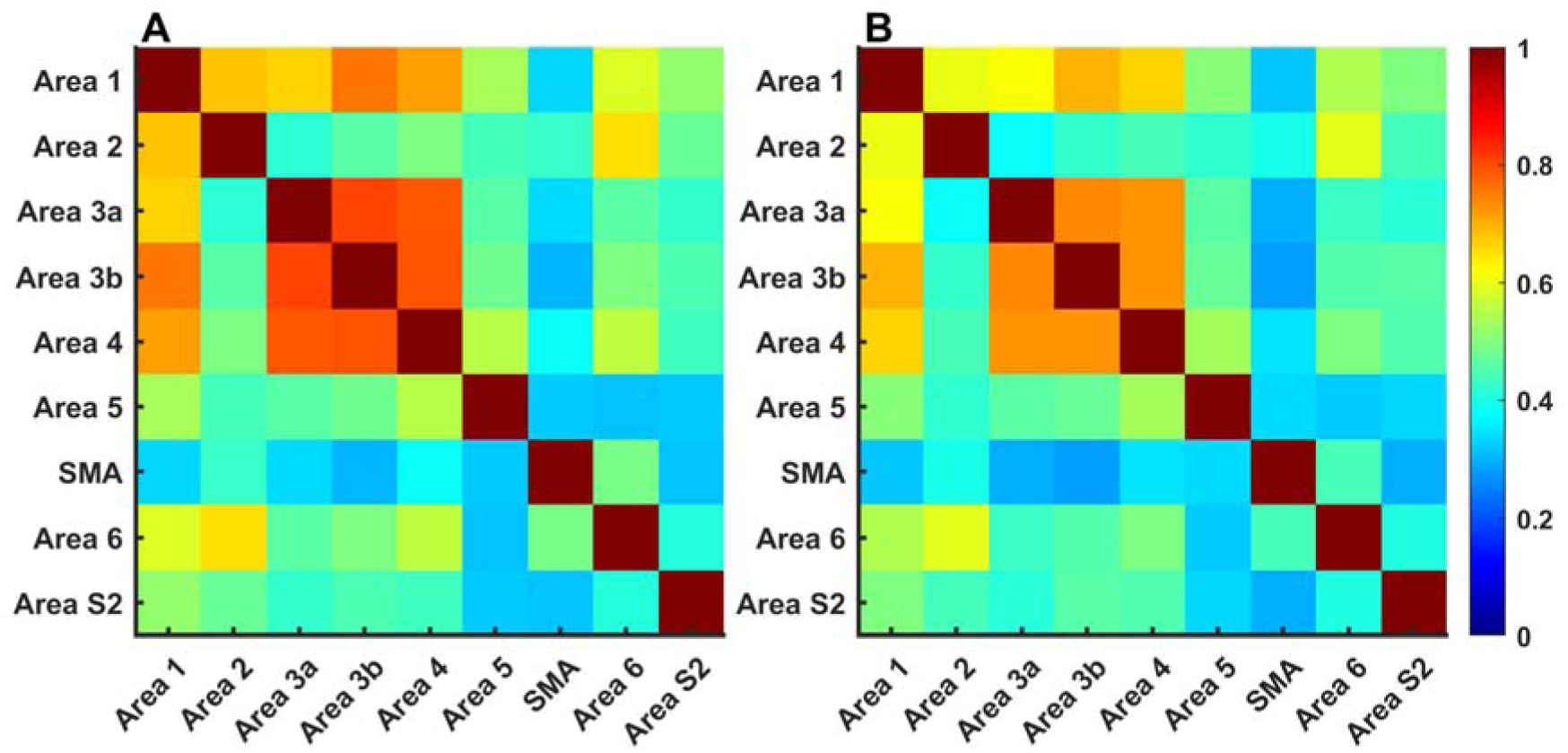
FC matrix for the somatomotor network for Dataset 2. (A) BPF and (B) EMD method. Conventions same as for Fig. 7.

Using the BPF method, the rsFC_BPF_ values between area 1 and other regions, i.e. area 2, area 3a, area 3b, area 5 and area S2 were relatively higher, ranging from 0.52 to 0.76, suggesting strong FC between these areas. However, cortical areas not known to have SC to area 1, such as area 4, SMA and area 6 show moderately high rsFC values (with area 4, 0.72; SMA, 0.33; area 6, 0.59). Here “high” value of FC is referred with respect to the threshold value (shown in Fig. 6(I)), i.e., the rsFC_BPF_ value exceeding the threshold value by a substantial margin. The presence of the FC value for regions that are structurally negative indicate that the BPF method overestimates the FC, i.e., the regions appear to be functionally connected, but there are no structural pathways to support that connection (Felleman and Van Essen, 1991).

In contrast, EMD reveals FC between area 1 and area 2, area 3a, area 3b, area 5 and S2 with connectivity values in the range from 0.50 to 0.70, which are higher than the identified threshold value (Fig. 6 (J), thereby signifying positive FC for the structurally positive areas. For the structurally negative areas, the rsFC _EMD-HT_ values were 0.67 for area 4, 0.31 for SMA, and 0.50 for area 6. Notably the FC between the area 1-SMA is below the threshold. Thus, EMD method is better at not showing the FC between the areas that donot have SC.

For SMA also the BPF method overestimates the rsFC_BPF_. It showed FC for the structurally unsupported regions such as area 3a (0.34) and area 1 (0.33), both of which marginally exceed the optimal threshold identified in Fig. 6 (I). However, EMD selectively retains only structurally supported connections. For example, connections of SMA to areas 2, 4, 5, and 6 are preserved with rsFC _EMD-HT_ with values ranging from 0.34 to 0.44. The weaker associations that fall below the threshold- such as SMA-area 1 (0.31), SMA-area 3a (0.30), SMA-area 3b (0.28), and SMA-area S2(0.30), were excluded. The region pairs that are excluded correspond to structurally negative pairs that are identified as functionally negative, indicating better specificity of the EMD based method.

An estimate of the functional structural correspondence using the confusion matrix derived measures is given in Table 3. These metrics are obtained by thresholding the rsFC_BPF_/ rsFC _EMD-HT_ values at 0.32/0.33. The analysis show that TP and TN are higher for the EMD method. The FN and FP are also lower for the EMD than the BPF method. Thus, the EMD is better at revealing functional connections that are structurally supported. Using EMD the accuracy improved from 63.89% to 69.44%, precision from 63.64% to 67.74%, specificity from 14.29% to 28.57% and F1-score from 76.36% to 79.34%.

**Table 3.** Confusion Matrix -derived performance metrics for Dataset 2.

| Method | TP | TN | FP | FN | Accuracy | Precision | Sensitivity | Specificity | F1-score |
| --- | --- | --- | --- | --- | --- | --- | --- | --- | --- |
| BPF | 42 | 4 | 24 | 2 | 63.89% | 63.64% | 95.45% | 14.29% | 76.36% |

| Method |  |  |  |  |  |  |  |  |  |
| --- | --- | --- | --- | --- | --- | --- | --- | --- | --- |
| EMD Method | 42 | 8 | 20 | 2 | 69.44% | 67.74% | 95.45% | 28.57% | 79.34% |

TP, true positive; TN, true negative; FN, false negative; FP, false positive.

**Table 4.**
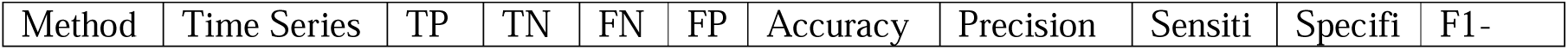
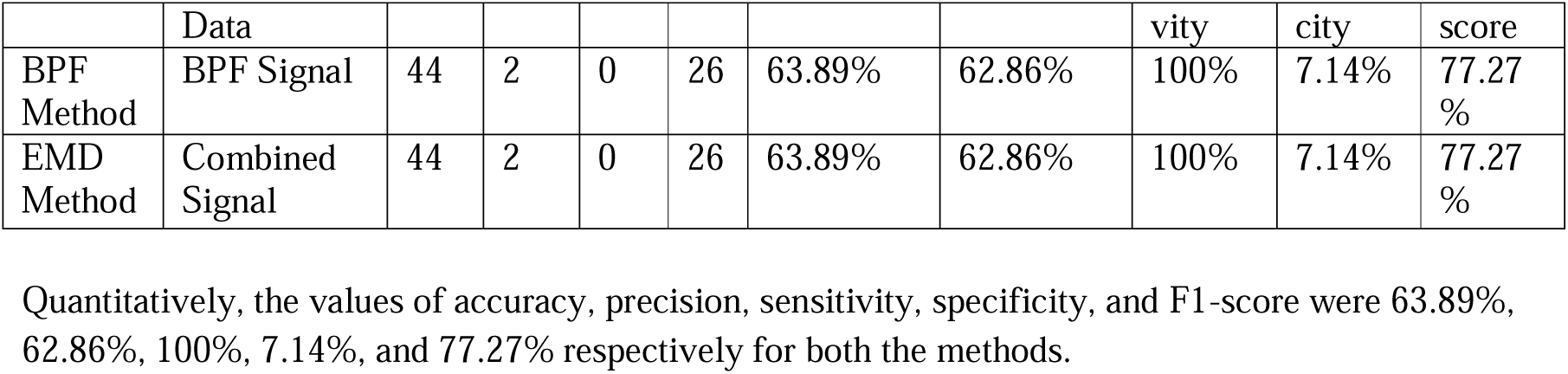
Confusion Matrix-derived performance metrics for Dataset 3.

| Method | Time Series | TP | TN | FN | FP | Accuracy | Precision | Sensiti | Specifi | F1- |
| --- | --- | --- | --- | --- | --- | --- | --- | --- | --- | --- |

|  | Data |  |  |  |  |  |  | vity | city | score |
| --- | --- | --- | --- | --- | --- | --- | --- | --- | --- | --- |
| BPF Method | BPF Signal | 44 | 2 | 0 | 26 | 63.89% | 62.86% | 100% | 7.14% | 77.27% |
| EMD Method | Combined Signal | 44 | 2 | 0 | 26 | 63.89% | 62.86% | 100% | 7.14% | 77.27% |

The structural-functional correspondence for Dataset 2 using both BPF and the EMD methods is shown in Fig. 10. For the BPF method, there were 12 functional-only (not revealed in SC) and 1 structural-only mismatches (not revealed in FC). With the EMD-based method, these were reduced to 10 and 1, respectively. Thus, the rsFC_EMD-HT_ computed via the EMD aligns more closely with the underlying structure. Moreover, using the BPF method the nodes with a greater degree centrality were involved in the functional-only mismatches, suggesting that the apparent functionally important hubs reflect structurally unsupported connectivity (Fig. 10). In contrast, the EMD not only reduced these mismatches but also preserved the centrality of the key hubs, suggesting that the thus identified functional hubs are more structurally grounded.

**Fig. 10.**
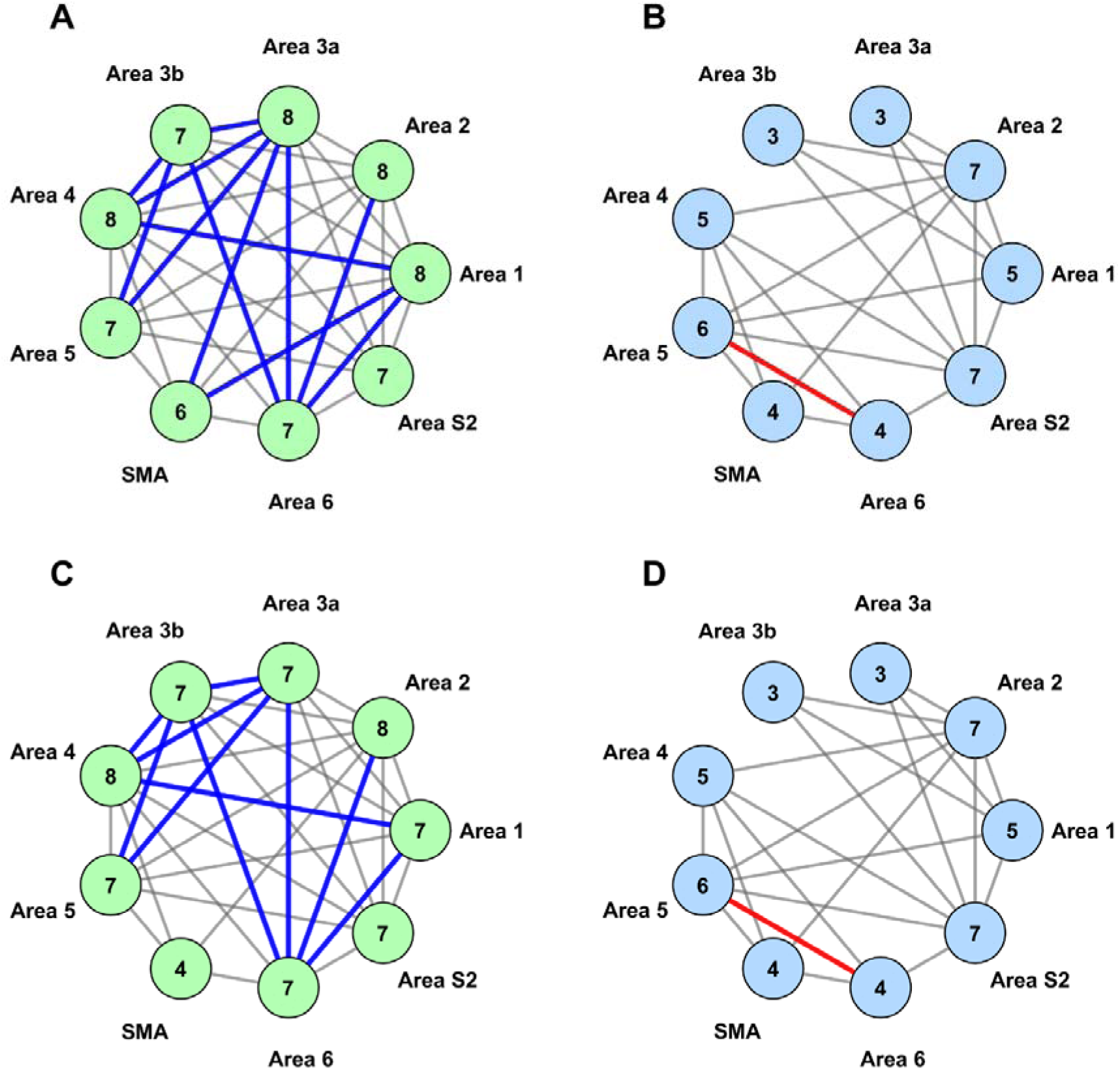
Structural functional mismatch (A) BPF method functional Connections not revealed in SC, (B) BPF method structural connections not revealed in FC, (C) EMD method functional connections not revealed in SC, (D) EMD method structural Connections not revealed in FC (Dataset 2), Legend description is same as in Figure 8

### 3.3 Dataset 3

In order to establish our conclusions, we used another resting-state fMRI dataset, which is available as part of the Amsterdam Open MRI collection. The Amsterdam PIOP2 dataset, which is available in pre-processed form has a same TR value of 2 seconds as our Dataset 1. We parcelled the pre-processed dataset using the Glasser atlas (Glasser et al., 2016) to extract the time series data from each voxel, which was then averaged across the ROI mask as outlined in the method section. A total of 10 subjects were taken for the analysis. The IMFs lying in the frequency range from 0.01 to 0.12 Hz were combined for analysis, and for thresholding the rsFC, values of 0.44 and 0.39 were taken.

The FC matrix for Dataset 3 is shown in Fig. 11. The results show that the EMD reveals FC that aligns closer with the SC. For example, SMA has known structural connections with cortical areas 2, 4, 5 and 6, while there is no SC with areas 1, 3a, 3b, and S2 (Felleman and Van Essen, 1991). When thresholding the functional connectogram at 0.44 to retain only the strongest connections, the BPF analysis includes regions that lack SC with SMA. Although it yields high rsFC_BPF_ for structurally connected pairs, such as 0.58 for SMA-area 2, 0.63 for SMA-area 4, 0.66 for SMA-area 5, and 0.74 for SMA-area 6, it also produced unexpectedly high values for structurally unconnected regions. For example, the rsFC_BPF_ values were 0.64 for SMA–area 1, 0.57 for SMA–area 3b, 0.47 for SMA–area 3a, and 0.41 for SMA-S2.

**Fig. 11.**
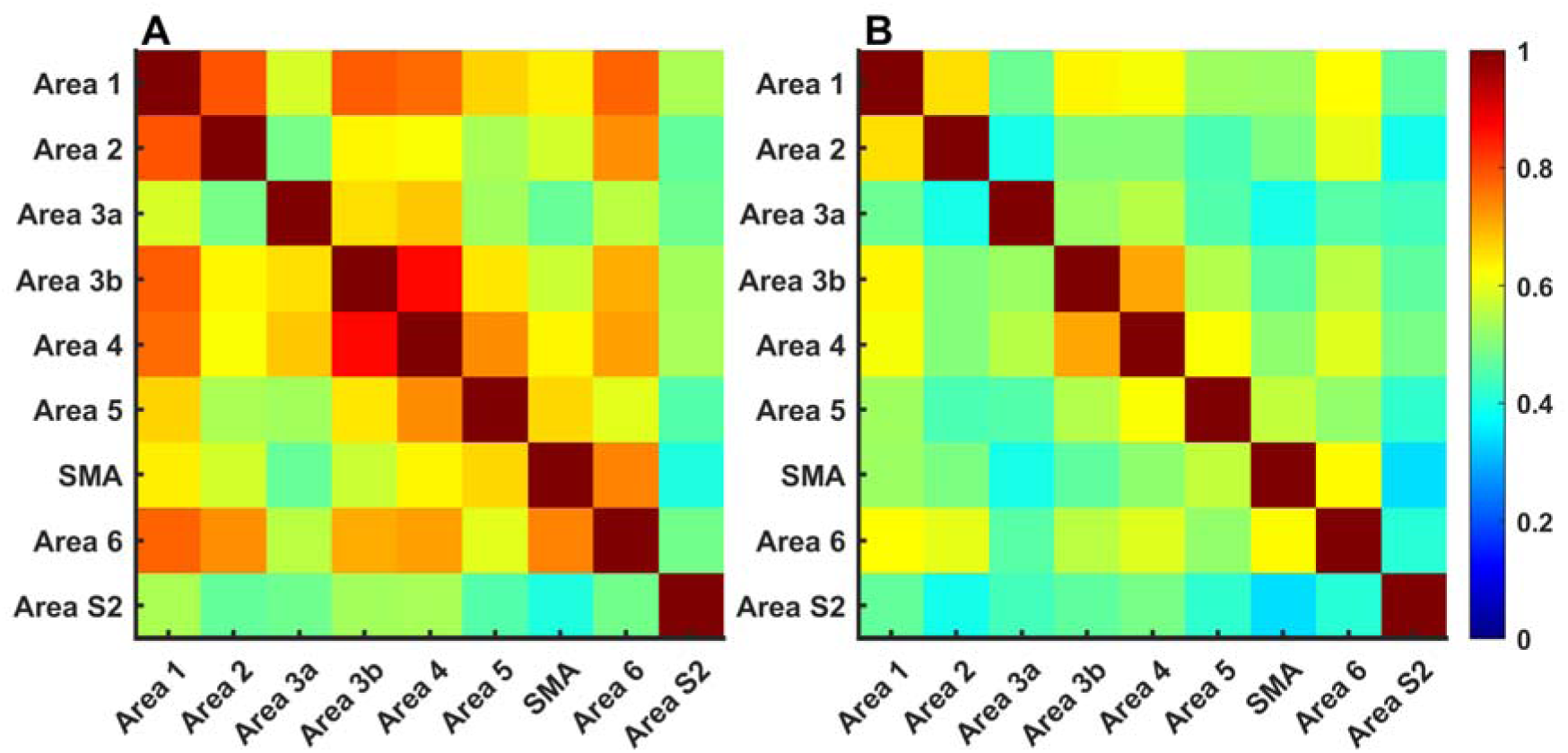
FC matrix for the somatomotor network for Dataset 3 using (A) BPF and (B) EMD methods. Conventions, same as for Fig. 7.

In contrast, EMD resulted in rsFC _EMD-HT_ values of 0.50 for SMA-area 2, 0.51 for SMA-area 4, 0.56 for SMA-area 5, and 0.63 for SMA-area 6. For regions with no structural connection to SMA, rsFC_EMD-HT_ values were lower than for rsFC_BPF._ For example, SMA-area 3a connectivity value drops from 0.47 in BPF to 0.40 using EMD, and SMA-area S2 from 0.41 to 0.34. Similarly, a decrease was observed for SMA-area 1, and SMA-area 3b, where the FC value dropped from 0.64 to 0.52 and from 0.57 to 0.47, respectively. Thus, EMD based FC more closely reflects the underlying SC than that revealed by the BPF method. Overall, both methods seem to indicate a strong rsFC for the region with known structural pathways, but the EMD method assigned a lower FC value to structurally unconnected regions.

Table 6 presents a comparative analysis of the performance measures derived from the confusion matrix for both the BPF and EMD-based methods. The metrics indicate that the classification performance using the EMD and BPF based method is same.

The interpretation is further supported by the mismatch analysis, which is shown in Fig. 12. It indicates the number of edges where the SC and FC’s are not aligned. In the functional-only mismatch (not revealed in SC), the BPF and EMD method both produced 13 mismatch edges. In the structural-only mismatches (not revealed in FC), both methods produced zero mismatched edges, indicating that neither method underrepresented the structurally valid connections.

**Fig. 12.**
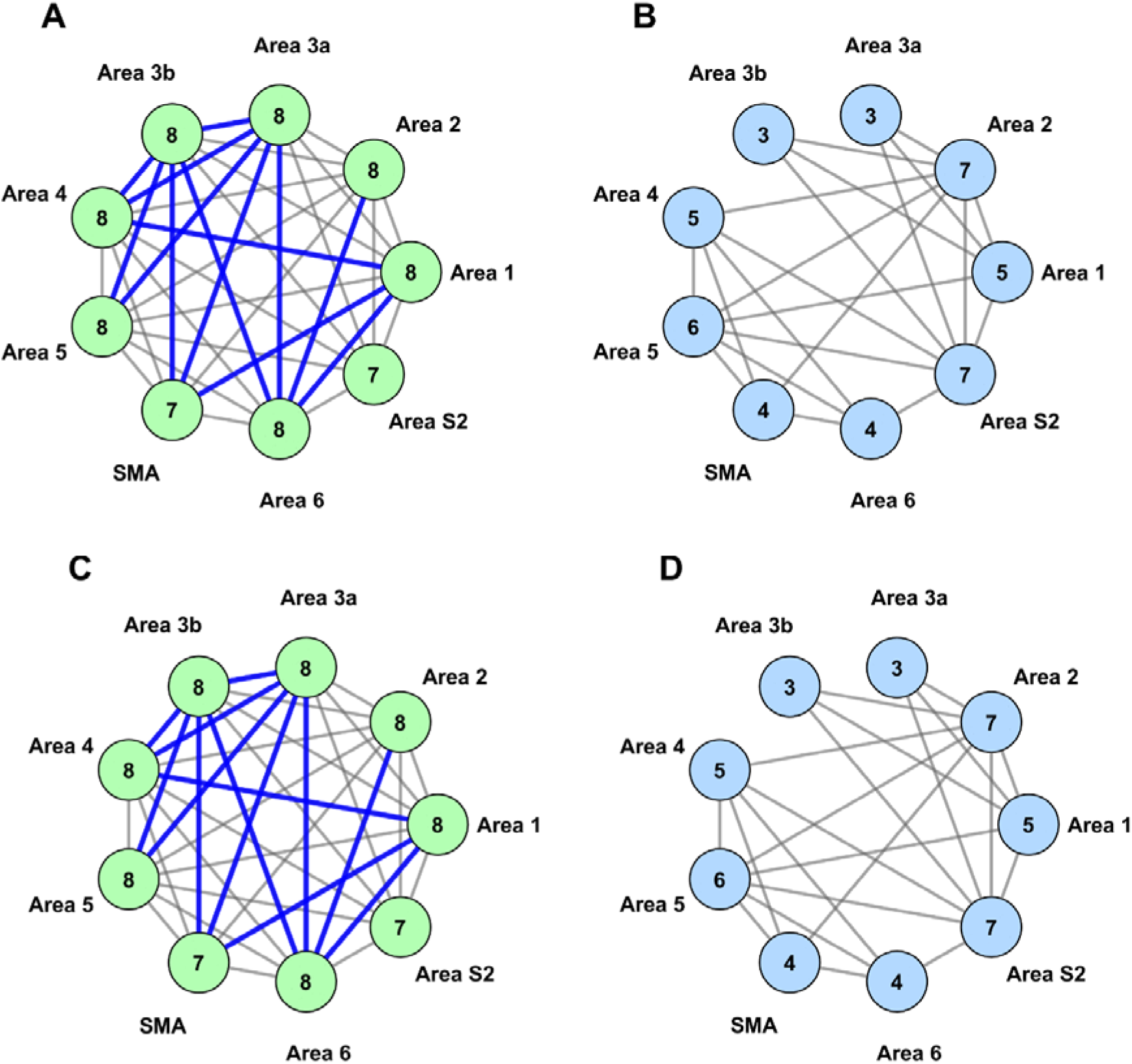
Structural functional mismatch (A) BPF method functional Connections not revealed in SC, (B) BPF method structural connections not revealed in FC, (C) EMD method functional connections not revealed in SC, (D) EMD method structural Connections not revealed in FC (Dataset 3), Legend description is same as in Figure 8.

## 4. DISCUSSION

Resting state functional networks in the brain capture the brain dynamics arising from the temporally coordinated fluctuations when it is not actively performing any task. Here, we have used Empirical Mode Decomposition and Hilbert Transformation (EMD-HT) to determine the resting state functional connectivity of the somatomotor network from the BOLD time series data and compared it with the more commonly used simple Band-Pass Filter (BPF) method. We show that EMD-HT improves the correspondence between the derived functional connectivity (FC) and the known structural connectivity (SC).

### 4.1 Functional connectivity and the underlying anatomical connectivity

The resting state functional connectivity is largely dependent on the underlying anatomical architecture of the brain. Macaque cortex is organized in a hierarchical fashion with projections (both feedforward and feed backward) linking the sensory, motor and association regions in a systematic order Felleman and Van Essen (Felleman and Van Essen, 1991). The structural connectivity provides a scaffold that shapes the cortical dynamics, such that the stronger structural links tends to have higher probability of synchronized activity even in the resting state (Honey et al., 2009). By decomposing the BOLD signal into physiologically meaningful IMF’s, rsFC_EMD-HT_ computes FC using the signal generated from the combination of these IMF’s, highlighting the patterns that align closely with the hierarchical organized cortical networks.

### 4.2 Function connectivity emerges even in the absence of direct structural connectivity

Functional connectivity is not constrained by direct structural connectivity between the brain regions(Raichle, 2011). FC between brain areas is observed even in absence of monosynaptic connections (Vincent et al., 2007).

Our results also reveal that the FC exists for regions that lack the direct structural connections. These correlations might have arisen due to indirect structural linkages. For example, we found a functional association between area 1-area 4, area 1-area 6, area 3a-area 3b, area 3a-area 4, area 3a-area 5, area 3b-area 4, area 3b-area 5 and area 3b-area 6 consistently across all three datasets.

### 4.3 Use of EMD for rs-fMRI analysis

EMD is often used for the analysis of non-linear and non-stationary signals (Huang et al., 1998). It has also been used for rsfMRI analysis (Zhang et al., 2015) (Szakál, 2016) (Yuen et al., 2019).

Zhang et al. (Zhang et al., 2015) proposed Multivariate EMD (MEMD) for resting state fMRI decomposition. Using resting state fMRI data of 5 minutes (TR=2 seconds) from 25 subjects, they determined. DMN network using seed-based analysis. They show that MEMD can adaptively decompose the fMRI time series data into IMFs. These IMFs reveal distinct patterns of the DMN connectivity, with connectivity values and the organization varying across IMFs. The comparison across IMFs reveals that the intermediate IMFs encompassing the frequency range from 0.01 to 0.08 exhibit strong within DMN functional connectivity that is consistent across subjects.

Vakarov et al. (Szakál, 2016) used EMD to study the resting state FC patterns of two brain regions, i.e., the visual cortex and the motor cortex. They identified the strongest temporal correlations between the homologous interhemispheric areas (within the same functional zone) and the visual areas. The weakest correlations were seen between the inter-zone pairs, such as motor-occipital connections. These findings were consistent among the IMFs, with the 2^nd^ IMF showing the strongest connectivity value. Also, the authors concluded that EMD is a useful method for extracting meaningful excerpts in the low-frequency range from resting-state fMRI data. Yuen et al. (Yuen et al., 2019) used VMD to decompose each voxel in the functional data into IMFs that were clustered around 0.22 Hz, 0.15 Hz, 0.080 Hz, and 0.028 Hz, corresponding to IMF1, IMF2, IMF3, and IMF4. These frequency clusters were found to be reproducible irrespective of the functional data sampling rate. They further show that the brain network topology can be successfully constructed using IMF 3 (dynamic range, 0.063–0.098) and IMF4 (dynamic range, 0.021–0.036 Hz),and found that the correlation values obtained using IMF3 and IMF4 were lower than that obtained from the band-pass method.

Here we determined if FC is determined using EMD-HT, will it correlate better with the known SC of the somatomotor network.

### 4.4 EMD-derived connectivity relates better to the structural connectivity

Our results indicate that EMD-derived connectivity relates better to the SC particularly in datasets with lower TR. The advantage arises because EMD effectively decomposed the rs-fMRI signal into frequency components characterising somatomotor network dynamics, while instantaneous frequency provided finer details of the IMF’s oscillatory behaviour at each time point.

Compared to conventional BPF methods, the EMD method better characterised the temporal fluctuations in brain activity. The improved alignment between structural and FC observed in EMD-HT arises from its data driven ability and the adaptivity in capturing the nonstationary fMRI signal characteristics. Unlike the BPF, which applies a fixed frequency window, under the assumption that the frequencies remain constant over time, EMD decomposes the ROI time series into IMF’s directly from the data itself. Furthermore, applying HT to these IMFs provides an estimate of how the instantaneous frequency and amplitude vary over time, thereby capturing the evolution of region-specific neural dynamics and accommodating the inherent dynamic nature of the BOLD signal.

In our results, these characteristics enabled EMD to preserve time-varying interactions between the resting state networks which might have been blurred when analysed using the BPF. Additionally, the rapid shift in the oscillatory activity was measured by the instantaneous frequency which gave a fine-grained representation of the network dynamics.

The EMD derived FC exhibited patterns that closely paralleled the known structural pathways. In contrast the BPF method did not account for the temporal fluctuations in the neural oscillations leading to the potential loss of the information when the frequencies drift a little outside the pass band. Additionally, the BPF method can introduce phase distortion, temporal blurring, and edge effects, which may obscure the true timings and the amplitude relationship between the time series data from different ROI’s thereby reducing the specificity of the connectivity estimates and their correspondence with the SC. The above findings underscore that EMD based analysis is a powerful tool for investigating the dynamic nature of the FC and the structure-function coupling in the human brain.

### 4.5 Performance comparison across different TR’s

It has been suggested that there are correlations between distant brain regions at higher frequencies, which can be estimated using the low-TR fMRI(Kalcher et al., 2014). The loss in the image resolution is often outweighed by increase in temporal resolution, as faster sampling provide a detailed view of the underlying dynamics. Consistent with this idea, we compared the results across the different TRs, where we observed that the accuracy is higher for the dataset having a lower TR and a larger number of time points (TR=0.72, number of time points=1200, accuracy=69.44%) than for the dataset having a higher TR and lower number of time points (TR=2, number of time points=205/240, accuracy =58.33%/63.89%). This is possibly because at lower values of TR, we have a higher temporal resolution, and as a result, subtle and transient fluctuations are easily captured. These more detailed dynamics enable the FC estimates to be more consistent with the brain structural architecture.

Additionally, similar results were seen for BPF and the EMD-HT method at TR=2 because at higher values of TR the temporal resolution is low, limiting the ability to capture fast fluctuations.

## 5. Conclusion

We evaluated if the resting state functional connectivity determined using EMD and Hilbert transformation (HT) has better correlation with the structural connectivity as compared to more commonly used methods. For this, we experimented on resting-state functional data of ten subjects each from three different datasets. The rs-fMRI was pre-processed, and thereafter, time series data were extracted from different brain regions using the Glasser parcellation. We thereafter decomposed the time series using EMD and further segregated the IMF’s holding the frequency of interest using HT. Mostly, IMF2, IMF3, and IMF4 were capturing the frequency band of interest in all the subjects. We reconstructed the signal using IMF2-4 and then estimated the functional connectivity. Our analysis indicates that the dynamics of the non-linear and non-stationary BOLD signal were better captured using the data-driven approach, as we observe a higher value for the structural-functional correspondence. We also validated our method on available standard parcelled/pre-processed datasets, where we also observed a similar trend. For all these datasets, the outcomes are better aligned with the structural basis.

## Acknowledgement

We are thankful to John Thomas for acquistion of Dataset 1.

## Notes

### Competing Interest Statement

The authors have declared no competing interest.

### Summary of Updates

The title has been revised and 2-3 lines in the abstract and introduction are revised.

